# A multigenerational Turing model reproduces transgressive petal spot phenotypes in hybrid *Mimulus*

**DOI:** 10.1101/2023.08.10.552669

**Authors:** Emily S. G. Simmons, Arielle M. Cooley, Joshua R. Puzey, Gregory D. Conradi Smith

## Abstract

The origin of phenotypic novelty is a perennial question of genetics and evolution. To date, few studies of biological pattern formation specifically address multi-generational aspects of inheritance and phenotypic novelty. For quantitative traits influenced by many segregating alleles, offspring phenotypes are often intermediate to parental values. In other cases, offspring phenotypes can be transgressive to parental values. For example, in the model organism *Mimulus* (monkeyflower), the offspring of parents with solid-colored petals exhibit novel spotted petal phenotypes. These patterns are controlled by an activator-inhibitor gene regulatory network with a small number of loci. Here we develop and analyze a model of hybridization and pattern formation that accounts for inheritance in diploid gene regulatory networks composed of either homozygous or heterozygous alleles. We find that the resulting model of multi-generational Turing-type pattern formation can reproduce the transgressive petal phenotypes observed in *Mimulus*. The model gives insight into how non-patterned parent phenotypes can yield phenotypically transgressive, patterned offspring, aiding in the development of empirically testable hypotheses.

## 1 Introduction

The origin of phenotypic novelty is a perennial question of evolutionary genetics. In answering this question, evolutionary biologists often focus on intergenerational changes over long time scales. However, it is well known that a dramatic amount of phenotypic variability can arise within a single generation. In plant biology, for example, breeding divergent varieties produces offspring whose yield is significantly greater than the yield of either parent, a phenomenon known as heterosis (i.e., hybrid vigor). In heterosis, the interaction between alleles from different parents somehow produces an emergent phenotype. This paper is a theoretical investigation of this source of phenotypic novelty.

Previous work has demonstrated that crosses between different *Mimulus* species with solid-colored petals can produce F1 offspring with spotted petals (P_A_ *×* P_B_ *→* F1). In particular, crosses of *M. luteus* var. *variegatus* (red) with *M. cupreus* (yellow or orange) produce F1 hybrids that are yellow with red spots (Fig. 1A, bottom two rows). The distribution of phenotypes in an F2 population generated from the selfing of F1 hybrid plants (F1 *×* F1 *→* F2) yields additional novelty [Cooley and Willis, 2009]. We observe a complex distribution of phenotypes with petals having a range of spot number, size, location, intensity, and color (Fig. 1B). What is the explanation for phenotypes changing so dramatically over just two generations? A working hypothesis must begin with the current understanding of the gene regulatory network controlling anthocyanin production, the red-purple pigment found in *Mimulus* petals.

**Figure 1:**
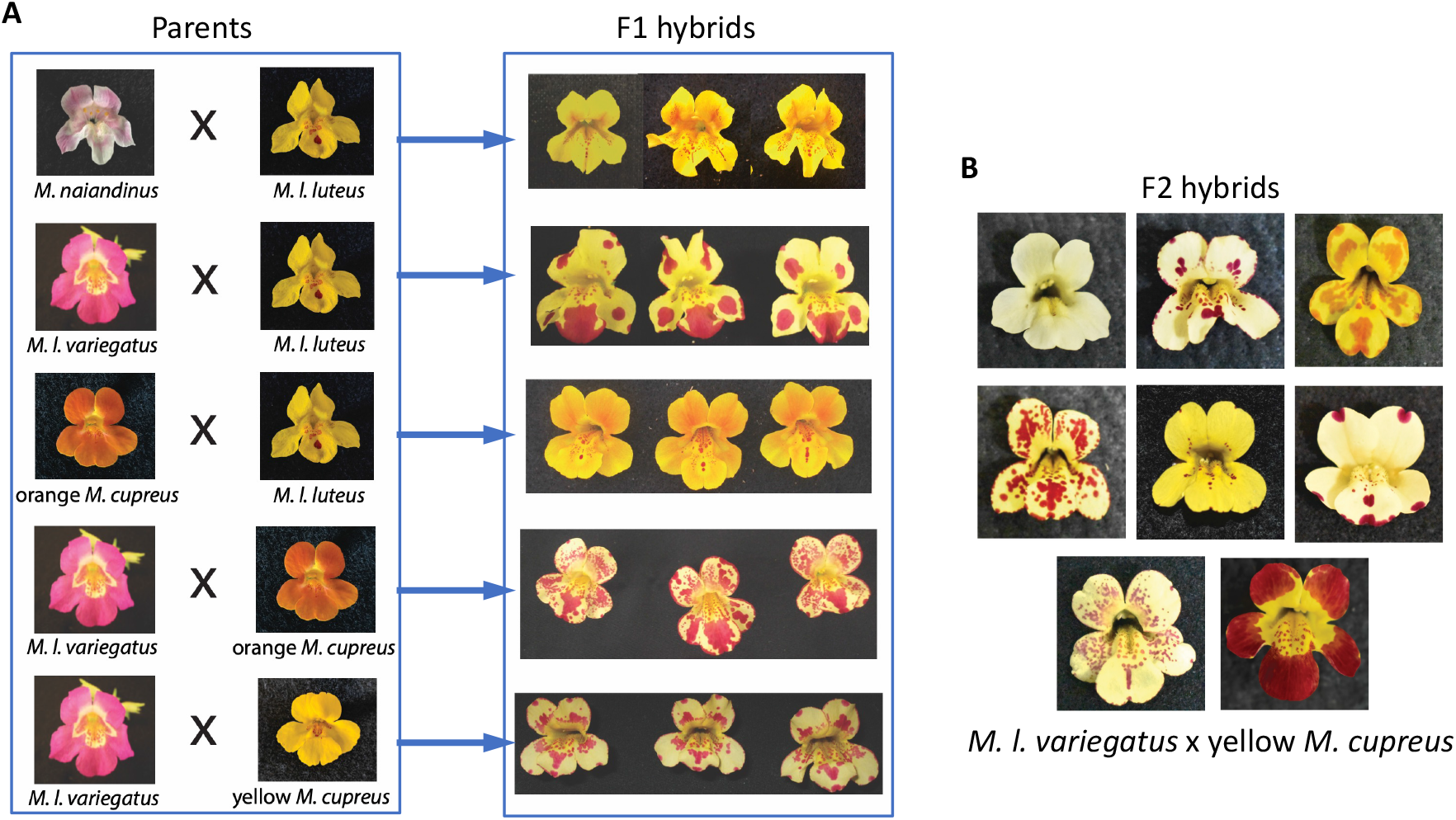
Transgressive petal phenotypes in crosses between *Mimulus* species. (A) Inbred (i.e., homozygous) parent flowers with unpatterned petal phenotypes produce reliable patterned phenotypes in F1 hybrids. (B) Self-pollination of F1 *M. l. variegatus × M. cupreus* hybrids yield anthocyanin patterns of varying size, density, and intensity of spots.

In monkeyflowers, the biochemical pathway that produces anthocyanin pigment is controlled by MYBs, a well-known transcription factor (TF) family (see Stracke et al., 2001). These include two R2R3-MYB TFs, NEGAN/MYB5 in the nectar guide and PELAN in the petal lobes, and one R3-MYB TF, red tongue (RTO) [Ding et al., 2020]. Two accessory proteins, a basic helix-loop-helix and a WD40 (MlANbHLH1 and MlWD40a, respectively), form a complex with the activator MYBs (called the MBW complex). The MBW complex has been widely hypothesized to operate as an activator within a reaction-diffusion mechanism, while the R3-MYB operates as an inhibitor [Bouyer et al., 2008; Ding et al., 2020; Ishida et al., 2008; Larkin et al., 1996; Pesch and Hülskamp, 2004]. Research on various MBW complexes has shown that the R3 MYBs (the inhibitors) move intercellularly (long-range inhibition), expanding away from the cells in which they are synthesized; R2R3 MYBs (the activators) do not (short-range activation) [Albert et al., 2014; Ding et al., 2020; Kurata et al., 2005]. These results are consistent with the requirements for Turing-type pattern formation [Meinhardt, 2012; Turing, 1990].

To confirm that a reaction-diffusion-mediated pattern formation is plausible in monkeyflowers, researchers performed genetic manipulation on a species whose wild type always produces spotted nectar guides. These results show that NEGAN RNAi knockouts exhibit nectar guides lacking anthocyanin (elimination of activator in nectar guide). Homozygous and heterozygous RTO mutants exhibit very high and intermediate amounts of anthocyanin compared to the wild type, respectively (suppression of inhibitor). Finally, WD40a knockout mutants were devoid of anthocyanin everywhere on the flower (elimination of activator throughout the flower) [Ding et al., 2020].

Based on these empirical results, Ding et al. [2020] hypothesized that a Turing-type reaction-diffusion mechanism mediates Mimulus petal patterning. They demonstrated that a simple two-variable (activator and inhibitor) reaction-diffusion system can mimic experimentally observed phenotypes (variation of spot size, elimination of spots). Their phenomenological model consists of the following two partial differential equations (PDEs),

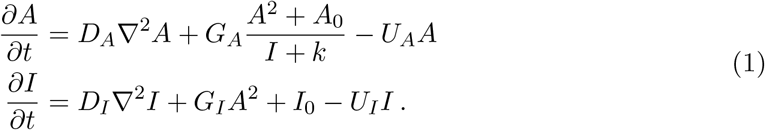

These PDEs account for the diffusion of activator and inhibitor and include nonlinear reaction terms for three of the four activator-inhibitor interactions typically found in Turing systems (no auto-inhibition) [Ding et al., 2020].

The Ding et al. model shows that activator-inhibitor reaction-diffusion equations can mimic the wild-type spotted nectar guide phenotype and experimental perturbations. This work is similar in spirit, but our focus is the appearance of novel (transgressive) patterned phenotypes in F1 hybrids and the variety of patterns found in the F2 generation. Section 2 shows how to model inheritance within the reaction-diffusion framework, beginning with reaction terms that instantiate a hypothesized mechanism of transcription factor binding and regulation. This enables the derivation of a reaction-diffusion system that accounts for diploidy—in particular, the effect of heterozygosity—on gene regulatory network function and pattern formation. Because genetic inheritance occurs by propagating parameters across simulated generations, we refer to our framework for modeling emerging petal patterns in hybrid *Mimulus* as a *multi-generational* Turing model.

## 2 Model formulation

The biochemical pathway that produces anthocyanin pigment in monkeyflowers involves activator and inhibitor transcription factor complexes. Our model formulation begins with a system of reaction-diffusion equations for an activator (*u*) and an inhibitor (*v*),

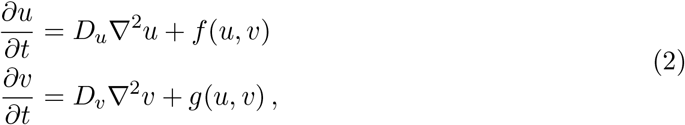

where *D*_*u*_ and *D*_*v*_ are diffusion coefficients. The reaction terms, *f* (*u, v*) and *g*(*u, v*), are

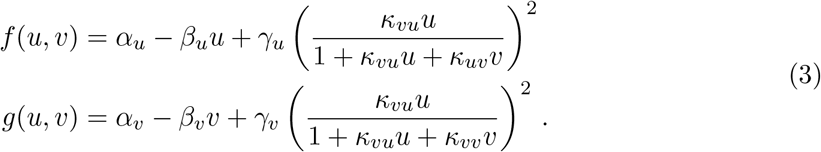

As illustrated in Fig. 2, these expressions represent an idealized gene regulatory network with the following properties. The activator *u* is produced at the baseline rate, *α*_*u*_, and degrades at a rate proportional to *u* (with a first-order rate constant *β*_*u*_). In addition, the production rate of *u* may be increased (by as much as *γ*_*u*_) through a Hill function that represents the competitive binding of activator *u* and inhibitor *v* to a pair of TF binding sites. For simplicity, these binding sites are assumed to be identical and independent. The inhibitor is modeled similarly, using the parameters *α*_*v*_, *β*_*v*_, and *γ*_*v*_.

**Figure 2:**
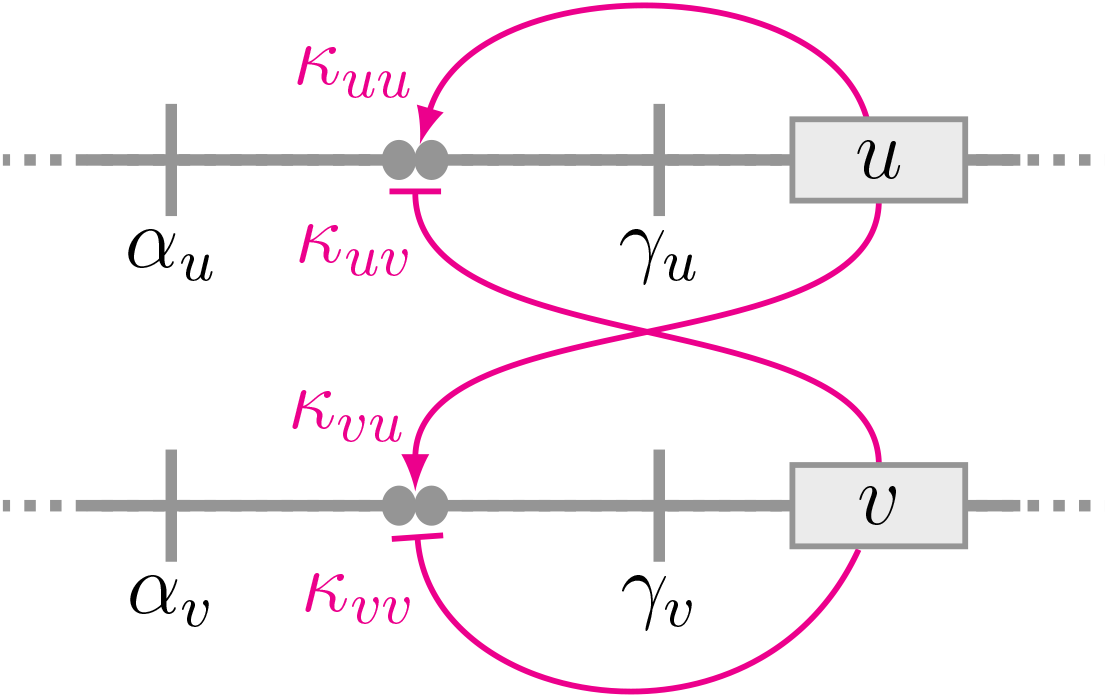
Schematic diagram of the activator-inhibitor gene regulatory network (haploid). The horizontal lines represent two unlinked genetic loci that code for transcription factors: an activator *u* and an inhibitor *v*. Filled circles represent transcription factor binding sites with affinities *κ*_*uu*_, *κ*_*uv*_, *κ*_*vu*_, and *κ*_*vv*_. The parameter *α*_*u*_ is the baseline production rate of activator *u*. The parameter *γ*_*u*_ is the increase in production rate that is regulated by the binding of the activator and the inhibitor (magenta curves). Similarly, *α*_*v*_ and *γ*_*v*_ are the baseline and regulatable production rate of inhibitor *v* (see Eq. 3).

The Hill functions occurring in Eq. 3 involve four equilibrium association constants: *κ*_*uu*_, *κ*_*uv*_, *κ*_*vu*_, and *κ*_*vv*_. In each case, the first subscript denotes the TF, *u* or *v*, whose production rate is being regulated. The second subscript denotes the TF that is binding. For example, *κ*_*uv*_ is the association constant for the inhibitor (*v*) binding to a site that regulates the activator production rate (see Fig. 2).

### 2.1 Accounting for diploidy

The reaction-diffusion system presented above (Eq. 2) represents two genetic loci (one for activator, one for inhibitor) known to exist on different chromosomes. To account for multi-generational aspects of *Mimulus* genetics, it is necessary to extend (duplicate) the model to represent two copies of each locus (one for each haploid genome). The resulting four-variable reaction-diffusion system takes the form

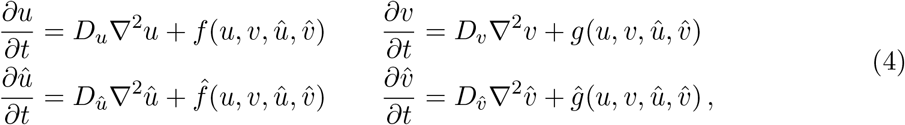

where *u* and *û* represent the (possibly distinct) gene products associated with the activator locus, and similarly for *v* and 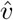 (see Fig. 3). The reaction terms appropriate for diploid *Mimulus* are an elaboration of those in Eq. 3,

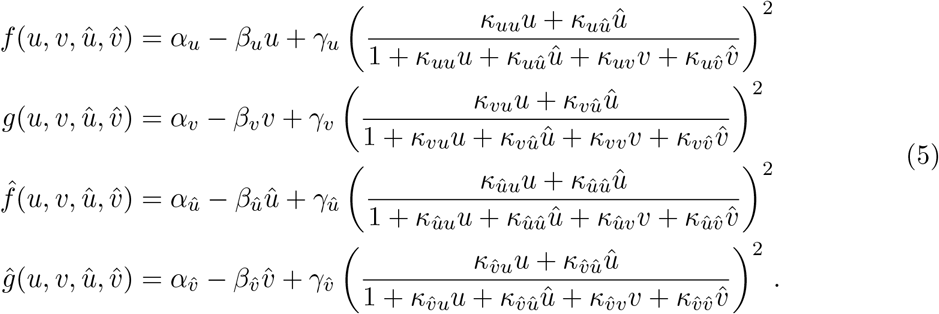

**Figure 3:**
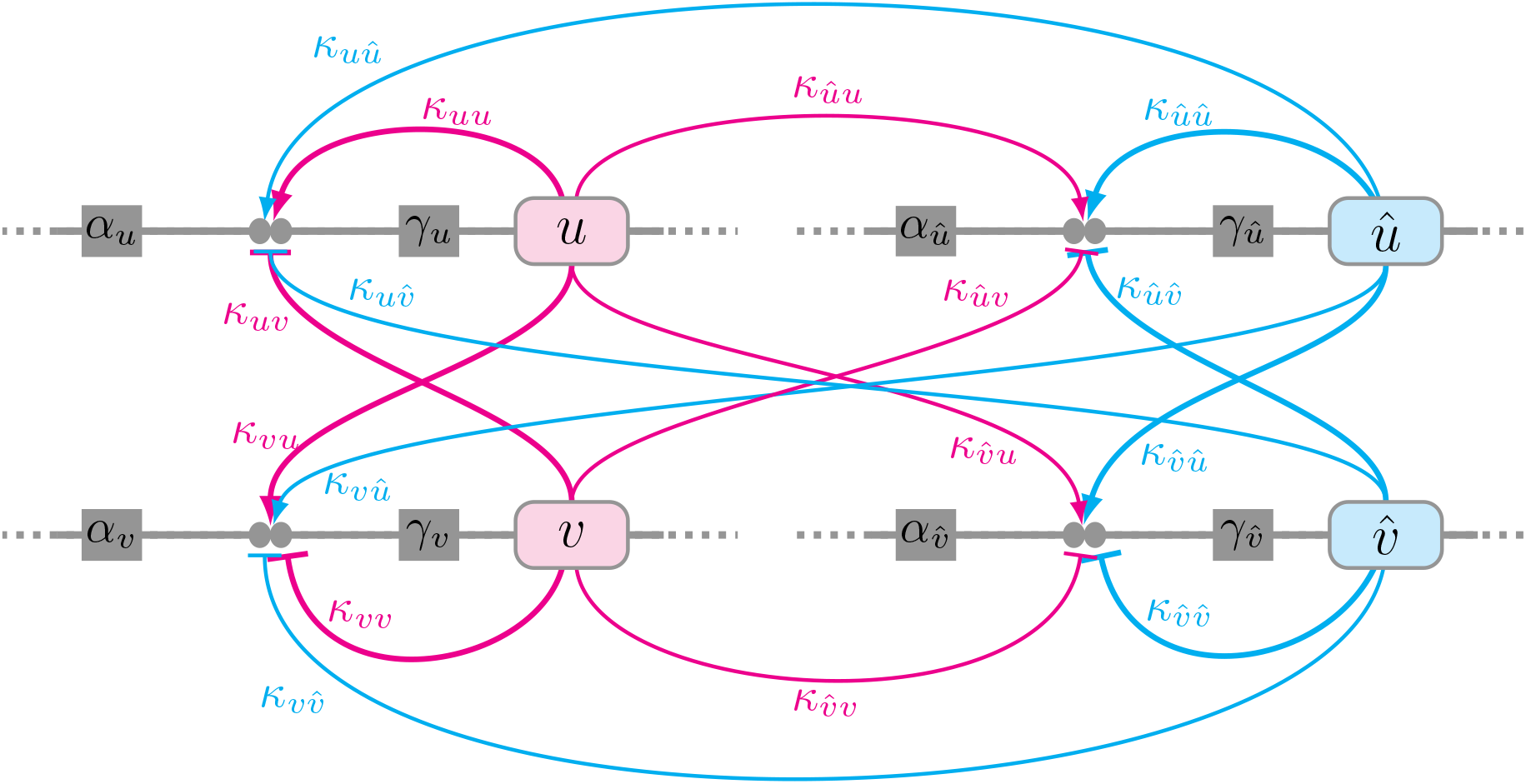
Schematic diagram of the *diploid* activator-inhibitor gene regulatory network. In essence, the diploid regulatory network is obtained by duplicating Fig. 2. There are four gene products, i.e., one activator and one inhibitor allele from each parent. The parental haplotype is distinguished by the presence or absence of a caret (*u* and *v* from parent 1, *û* and 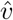 from parent 2). Filled circles represent transcription factor binding sites with affinities *κ*_*u\**_, *κ*_*û\**_, *κ*_*v\**_, and 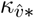 (where *\** is a placeholder for *u, v, û* and 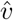). Thick and thin curves indicate an interaction between binding sites and transcription factors that are inherited from the same and opposite haplotypes, respectively.

The parameters *α*_*\**_, *β*_*\**_, and *γ*_*\**_ are the production rate, degradation rate, and maximum regulation rate of each of the four gene products (the subscript 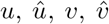). The four Hill functions in Eq. 5 include sixteen equilibrium association constants *κ*_*\*\**_. As in Eq. 3, the first subscript of these binding affinities denotes the TF (*u, û, v*, or 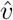) whose production rate is being regulated. The second subscript denotes which TF is binding (also, *u, û, v*, or 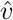 for a total of 16 parameters). For simplicity, our simulations assume that the rates of diffusion for both activator and inhibitor are independent of allele type (*D*_*û*_ = *D*_*u*_, 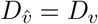). Consistent with Turing-type pattern formation, the diffusion coefficient for the inhibitor is assumed to be greater than activator *D*_*v*_ *> D*_*u*_).

### 2.2 Parameter assignment: homozygous parents and doubly heterozygous F1 hybrid

Consider heterozygous F1 offspring from the cross of two parents with distinct alleles for both activator and inhibitor (P_A_ *×* P_B_ *→* F1). Table 1 shows the assignment of rate constants 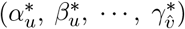 for the three diploid reaction-diffusion systems (Eq. 4) that model this cross. Superscripts indicate allele type, e.g., 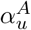 is the baseline production rate for the activator derived from parent A. The allele type chosen for each parameter follows Mendelian inheritance of the diploid genotypes for the activator (*U*_*AA*_, *U*_*AB*_, and *U*_*BB*_) and inhibitor (*V*_*AA*_, *V*_*AB*_, and *V*_*BB*_). Table 2 shows the inheritance of the association rate constants 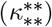. Here the interaction between the activator and inhibitor (both with two potentially different allele types) leads to the 16 possible values. The first subscript-superscript pair denotes the allele type of the TF (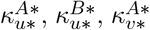, or 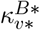) whose production rate is being regulated. The second subscript-superscript pair denotes the allele type of the TF that is binding 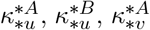, or 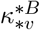. The F1 hybrid model uses all 16 binding constants. Each of the homozygous parents utilizes 4 binding constants (parent A highlighted pink, parent B highlighted cyan). That is, eight binding constants are only relevant for heterozygous *Mimulus*. Note: Although the genotype *U*_*BA*_ is not distinguishable from *U*_*AB*_, and similarly for *V*_*BA*_ and *V*_*AB*_. However, the binding constants 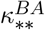 are distinct from 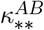. For example, the parameter 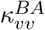 refers to the binding of the allele type A inhibitory transcription factor to the binding site associated with the regulation of type B inhibitor. The binding constant 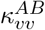 has a different interpretation and need not have the same value as 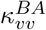.

**Table 1:**
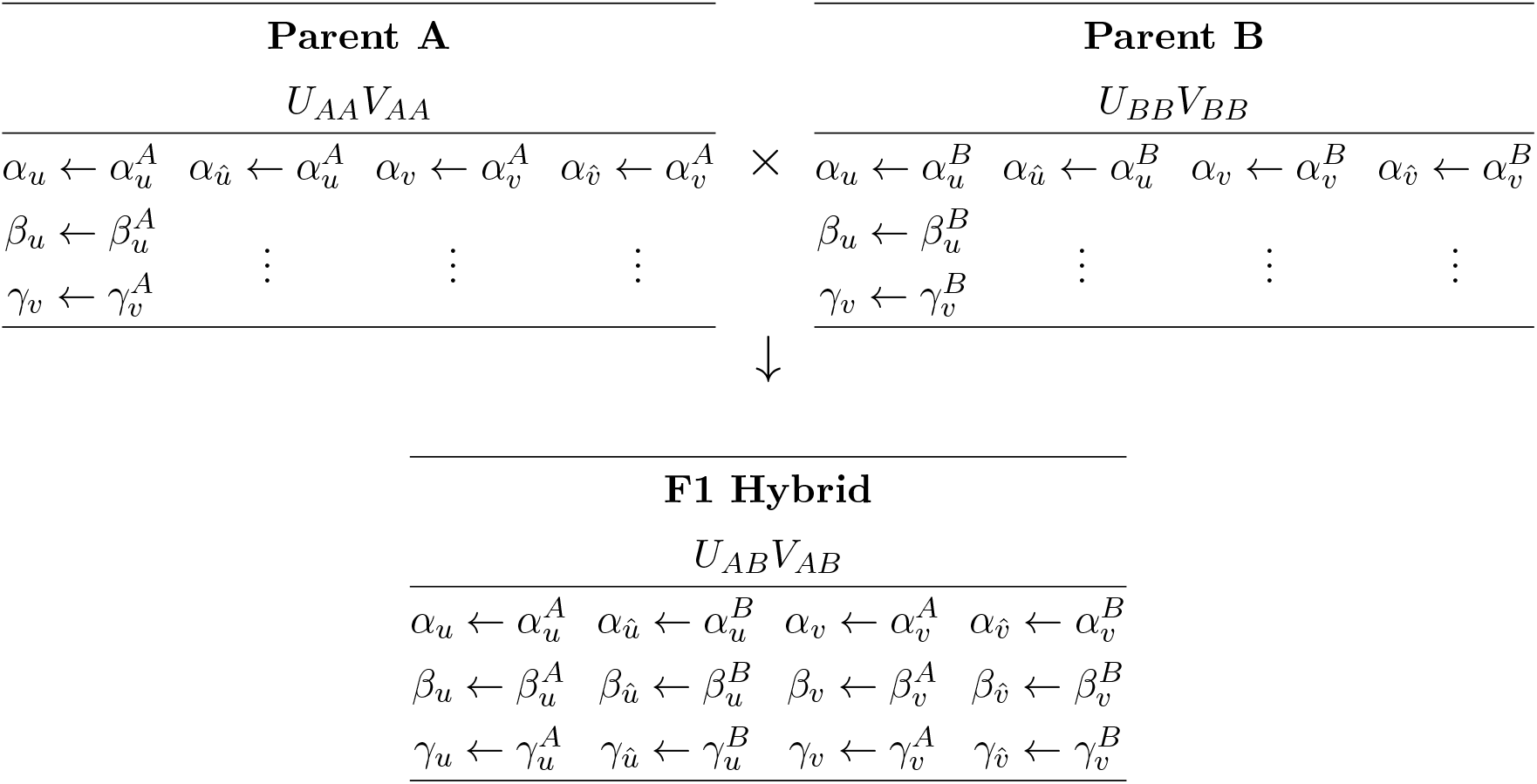
Assignments of rate constants for **P**_**A**_ *×* **P**_**B**_ *→* **F1 hybrid**.

**Table 2:**
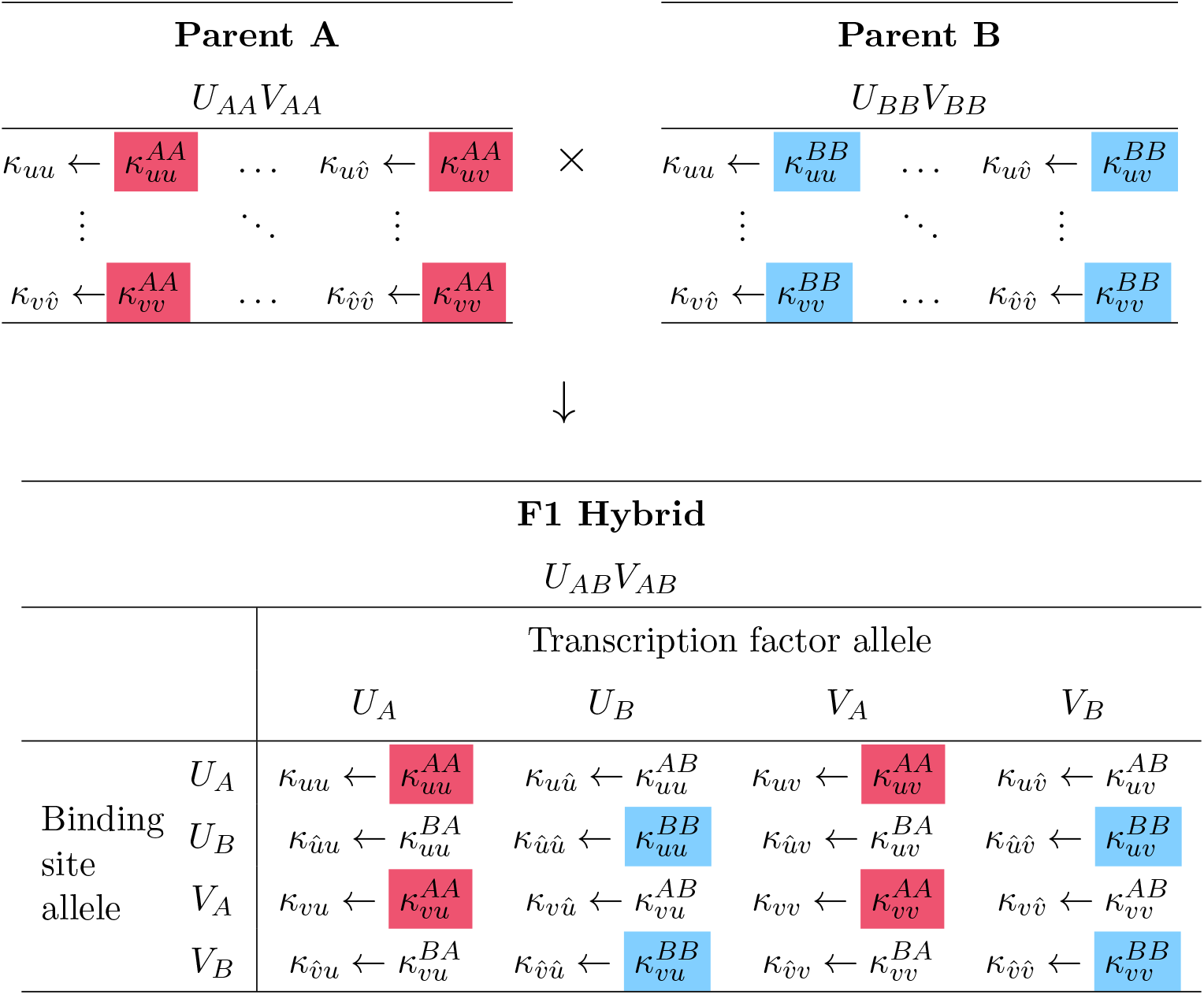
Assignment of binding constants for parents and F1 hybrid.

Using both Tables 1 and 2 and Eq. 4 we derive the reaction-diffusion system for the heterozygous F1 offspring of the P_A_ *×* P_B_ *→* F1 cross:

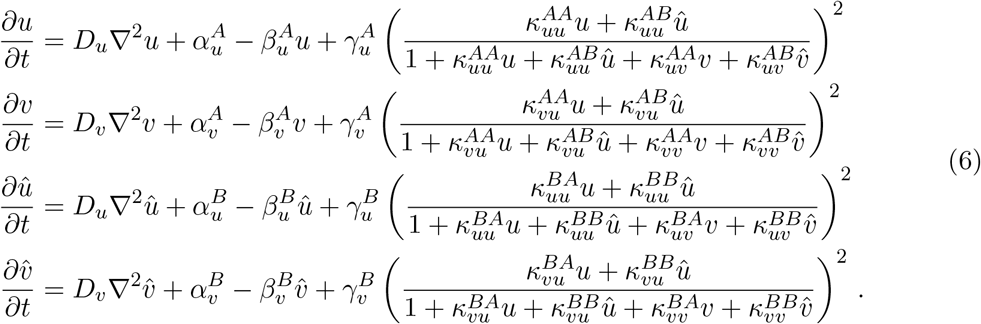

These equations can be compared and contrasted with the equations for parent A:

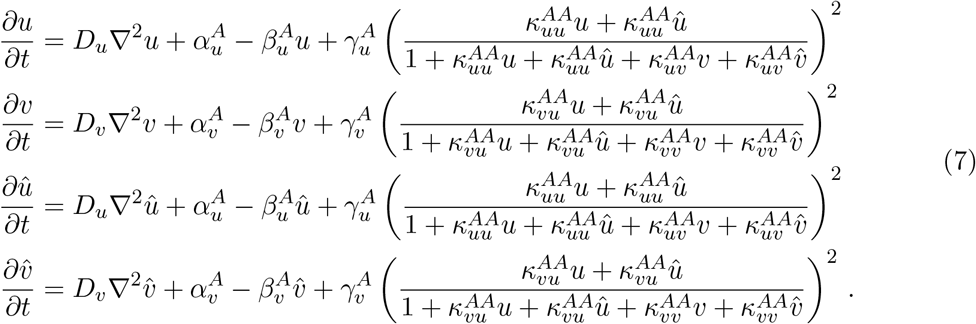

Because parent A is homozygous, the equations for *u* and *û* are identical, as are the equations for *v* and 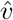. It is a simple matter to derive an equivalent reaction-diffusion system with one equation for each distinct gene product. Defining the total concentration of activator and inhibitor as 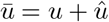 and 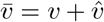, we arrive at the equivalent two-variable system,

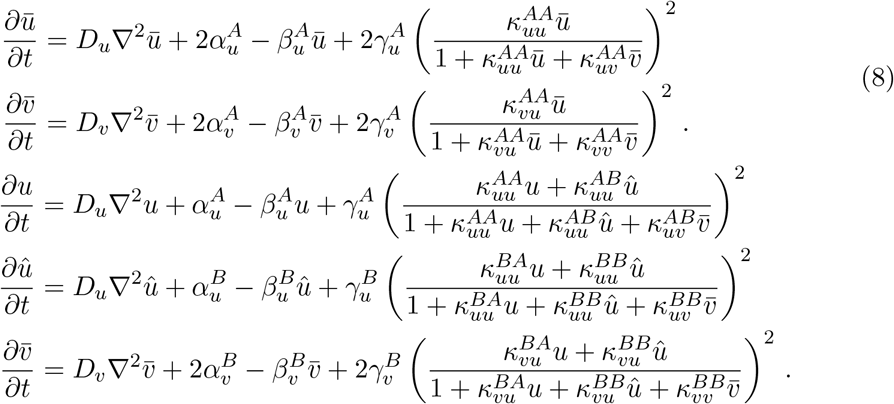

As expected, the zeroth-order rate constants (*α* and *γ*) are scaled by a factor of 2, while the first-order rate constants (*β*) are not. That is, a homozygous diploid model is equivalent to an appropriately scaled haploid model (compare Eq. 8 to Eqs. 2 and 3). The reaction-diffusion system for parent B (also homozygous) is identical to Eq. 7 with the replacement of *B* for *A*.

### 2.3 Inheritance of parameters in a simulated population of F2 hybrids

Hybrids of *M. cupreus* and *M. l. variegatus* show a striking distribution of phenotypes in the F2 generation (recall Fig. 1). In these experiments, the F2 hybrids are produced by selfing an F1 flower and growing up the progeny (F1 *×* F1 *→* F2). In our model, the distribution of genotypes in the simulated F2 generation is achieved by assigning parameters in Eq. 4 according to the Mendelian logic of the previous section.

Because the F1 hybrid is heterozygous for both activator and inhibitor (*U*_*AB*_*V*_*AB*_), the F2 hybrid population exhibits 9 distinct genotypes, including the doubly homozygous parental genotypes *U*_*AA*_*V*_*AA*_ and *U*_*BB*_*V*_*BB*_, the doubly heterozygous F1 hybrid genotype *U*_*AB*_*V*_*AB*_, and 6 genotypes that are unique to the F2 generation (*U*_*AA*_*V*_*AB*_, *U*_*AA*_*V*_*BB*_, *U*_*AB*_*V*_*AA*_, *U*_*AB*_*V*_*BB*_, *U*_*BB*_*V*_*AB*_, and *U*_*BB*_*V*_*AB*_). The parental and F1 cases were described in the previous section. For an example that is unique to the F2 hybrids, consider the *U*_*AB*_*V*_*BB*_ genotype, which is homozygous for activator and heterozygous for inhibitor. Simulations for this genotype use the rate and binding constants in Table 3.

**Table 3:** Assignment of rate and binding constants for the F2 hybrid genotype *U*_*AB*_*V*_*BB*_. The highlighted parameters differ from the doubly heterozygous F1 hybrid *U*_*AB*_*V*_*AB*_ (cf. Tables 1 and 2).

| **F2 Hybrid** |  |  |  |  |  |
| --- | --- | --- | --- | --- | --- |
| $U_{AB}V_{BB}$ | | | | | |
|  |  | Transcription factor allele |  |  |  |
| | | $U_A$ | $U_B$ | $V_B$ | $V_B$ |
| | | $\alpha_u \leftarrow \alpha_u^A$ | $\alpha_{\hat{u}} \leftarrow \alpha_u^B$ | $\alpha_v \leftarrow \alpha_v^B$ | $\alpha_{\hat{v}} \leftarrow \alpha_v^B$ |
| | | $\beta_u \leftarrow \beta_u^A$ | $\beta_{\hat{u}} \leftarrow \beta_u^B$ | $\beta_v \leftarrow \beta_v^B$ | $\beta_{\hat{v}} \leftarrow \beta_v^B$ |
| | | $\gamma_u \leftarrow \gamma_u^A$ | $\gamma_{\hat{u}} \leftarrow \gamma_u^B$ | $\gamma_v \leftarrow \gamma_v^B$ | $\gamma_{\hat{v}} \leftarrow \gamma_v^B$ |
| Binding<br>site<br>allele | $U_A$ | $\kappa_{uu} \leftarrow \kappa_{uu}^{AA}$ | $\kappa_{u\hat{u}} \leftarrow \kappa_{uu}^{AB}$ | $\kappa_{uv} \leftarrow \kappa_{uv}^{AB}$ | $\kappa_{u\hat{v}} \leftarrow \kappa_{uv}^{AB}$ |
| | $U_B$ | $\kappa_{\hat{u}u} \leftarrow \kappa_{uu}^{BA}$ | $\kappa_{\hat{u}\hat{u}} \leftarrow \kappa_{uu}^{BB}$ | $\kappa_{\hat{u}v} \leftarrow \kappa_{uv}^{BB}$ | $\kappa_{\hat{u}\hat{v}} \leftarrow \kappa_{uv}^{BB}$ |
| | $V_B$ | $\kappa_{vu} \leftarrow \kappa_{vu}^{BA}$ | $\kappa_{v\hat{u}} \leftarrow \kappa_{vu}^{BB}$ | $\kappa_{vv} \leftarrow \kappa_{vv}^{BB}$ | $\kappa_{v\hat{v}} \leftarrow \kappa_{vv}^{BB}$ |
| | $V_B$ | $\kappa_{\hat{v}u} \leftarrow \kappa_{vu}^{BA}$ | $\kappa_{\hat{v}\hat{u}} \leftarrow \kappa_{vu}^{BB}$ | $\kappa_{\hat{v}v} \leftarrow \kappa_{vv}^{BB}$ | $\kappa_{\hat{v}\hat{v}} \leftarrow \kappa_{vv}^{BB}$ |

Because this genotype is homozygous for the inhibitor (*V*_*BB*_) but not the activator (*U*_*AB*_), the corresponding reaction-diffusion system, after contraction, has three equations (*u, û* and 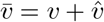):

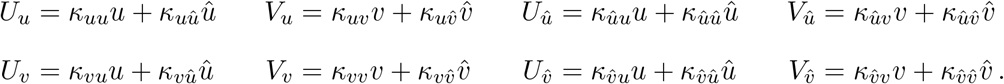

The reaction-diffusion equations for the remaining genotypes are derived similarly (see Appendix 4).

Note that the parameters used to simulate the doubly heterozygous F1 hybrid (*U*_*AB*_*V*_*AB*_) effectively define a given multi-generational simulation. Each of the other 8 genotypes (two parents, 6 unique F2 genotypes) use a subset of the F1 parameters. The F1 genotype equations (Eq. 6) use 16 association constants (see, e.g., Table 2). The doubly homozygous genotypes (*U*_*AA*_*V*_*AA*_, *U*_*AA*_*V*_*BB*_, *U*_*BB*_*V*_*AA*_, and *U*_*BB*_*V*_*BB*_) use 4 of these, while genotypes which are heterozygous at one locus and homozygous at the other use 9 (*U*_*AA*_*V*_*AB*_, *U*_*BB*_*V*_*AB*_, *U*_*AB*_*V*_*AA*_, and *U*_*AB*_*V*_*BB*_).

### 2.4 Parameters for transgressive hybrid phenotypes

Recall that a Turing instability for a two-variable activator-inhibitor system (e.g., Eqs. 2 and 3) requires linear stability of the reaction terms, *f* (*u, v*) and *g*(*u, v*). That is, the Jacobian defined by

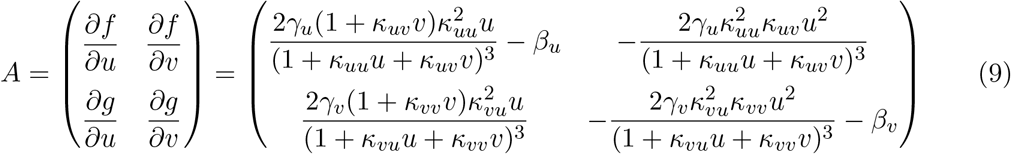

must be stable (both eigenvalues of *A*(*u*_*ss*_, *v*_*ss*_) must have negative real part) when evaluated at the steady state satisfying 0 = *f* (*u*_*ss*_, *v*_*ss*_) and 0 = *g*(*u*_*ss*_, *v*_*ss*_). Additionally, the matrix that arises from linearizing the full reaction-diffusion system,

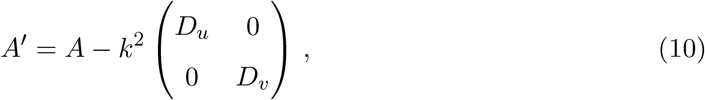

must be unstable (i.e., at least one eigenvalue has positive real part) for some spatial frequency *k*. It can be shown that this requires *d* = *D*_*v*_*/D*_*u*_ *>* 1 (see Murray, 2001, chap. 2).

For our model of patterning in diploid *Mimulus* (Eq. 4), the 4 *×* 4 Jacobian matrix is obtained by linearizing the reaction terms (Eq. 5). For compactness, these can be written as

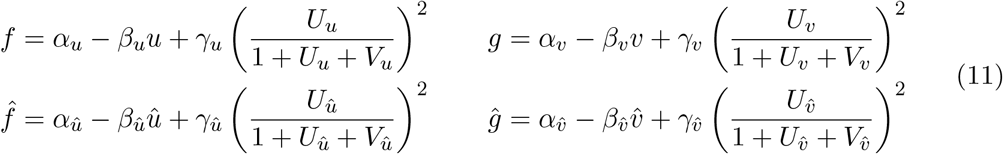

where

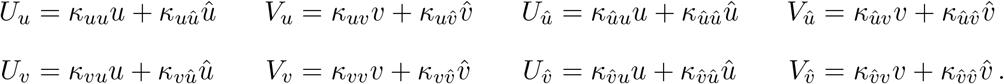

The Jacobian takes the form

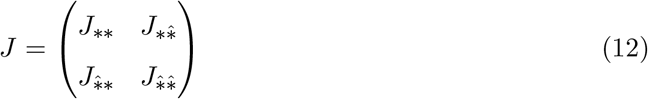

with diagonal blocks

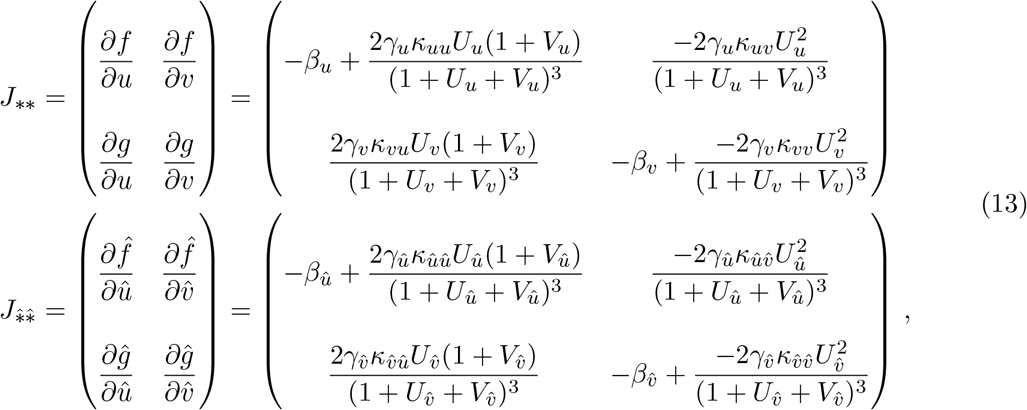

and off-diagonal blocks

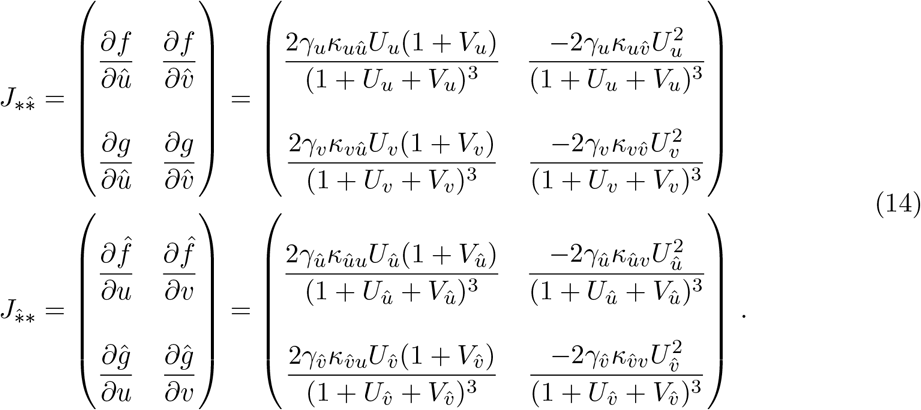

In these expressions, 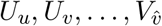 are evaluated at the steady state 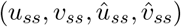 solving 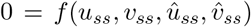, and similarly for *g*, 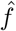, and *ĝ* (Eq. 11). A valid multi-generational parameter set (i.e., one resulting in a transgressive F1 phenotype) must yield three Jacobian matrices—*J* ^*A*^, *J* ^*B*^, and *J* ^*F* 1^—all of which are stable. Additionally, the matrix that results from linearizing the diploid reaction-diffusion system,

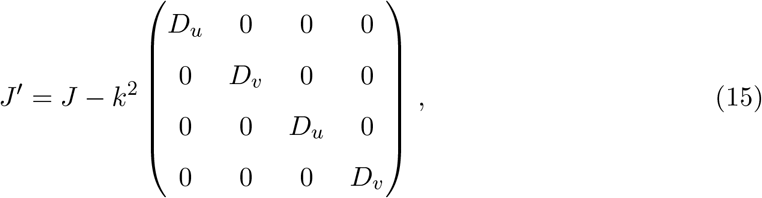

must be unstable for the F1 hybrid, yet stable for both parents.

When Eq. 15 is rewritten in terms of the relative diffusion coefficient, *d* = *D*_*v*_*/D*_*u*_, there exists a critical value of *d*, denoted *d*_*⋆*_, above which the matrix *J* ^*′*^ will be unstable for some spatial frequency *k* (a Turing bifurcation). Assume without loss of generality that 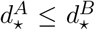. In that case, a valid multi-generational parameter set has critical diffusion ratios that satisfy 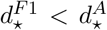. Choosing *d* = *D*_*u*_*/D*_*v*_ such that 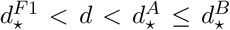, we obtain a multi-generational parameter set that results in Turing-stable parents and a Turing-unstable F1 hybrid (see Appendix B for further discussion).

## 3 Results

An intriguing observation of the experimental *Mimulus* system is that inbred (i.e., homozygous) parent flowers with unpatterned petal phenotypes produce reliable patterned phenotypes in F1 hybrids (recall Section 1). In particular, crosses between yellow morph *M. cupreus* and *M. l. variegatus* (both unpatterned) lead to spotted F1 hybrids (Fig. 1). To see if this phenomenon could be reproduced by our multi-generational model (Section 2), we sought parameter sets that yield this transgressive phenotype, that is, parameters for which Parents A and B are Turing stable, while the F1 hybrid is Turing unstable, as discussed above.

### 3.1 Unpatterned parents can produce patterned hybrid offspring

Fig. 4A shows a representative simulation in which parents A and B are unpatterned while the F1 hybrid is patterned. These simulations are performed using an alternating-direction-implicit Crank-Nicholson-like numerical scheme with no flux boundary conditions. As a proxy for the concentration of anthocyanin pigment, the steady-state value of total activator concentration (*u*+*û*) is shown in yellow-to-red pseudocolor. The total inhibitor concentration 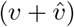 is not shown because it does not influence the visual appearance of *Mimulus* petals, but in these simulations, it is in phase with the activator.

**Figure 4:**
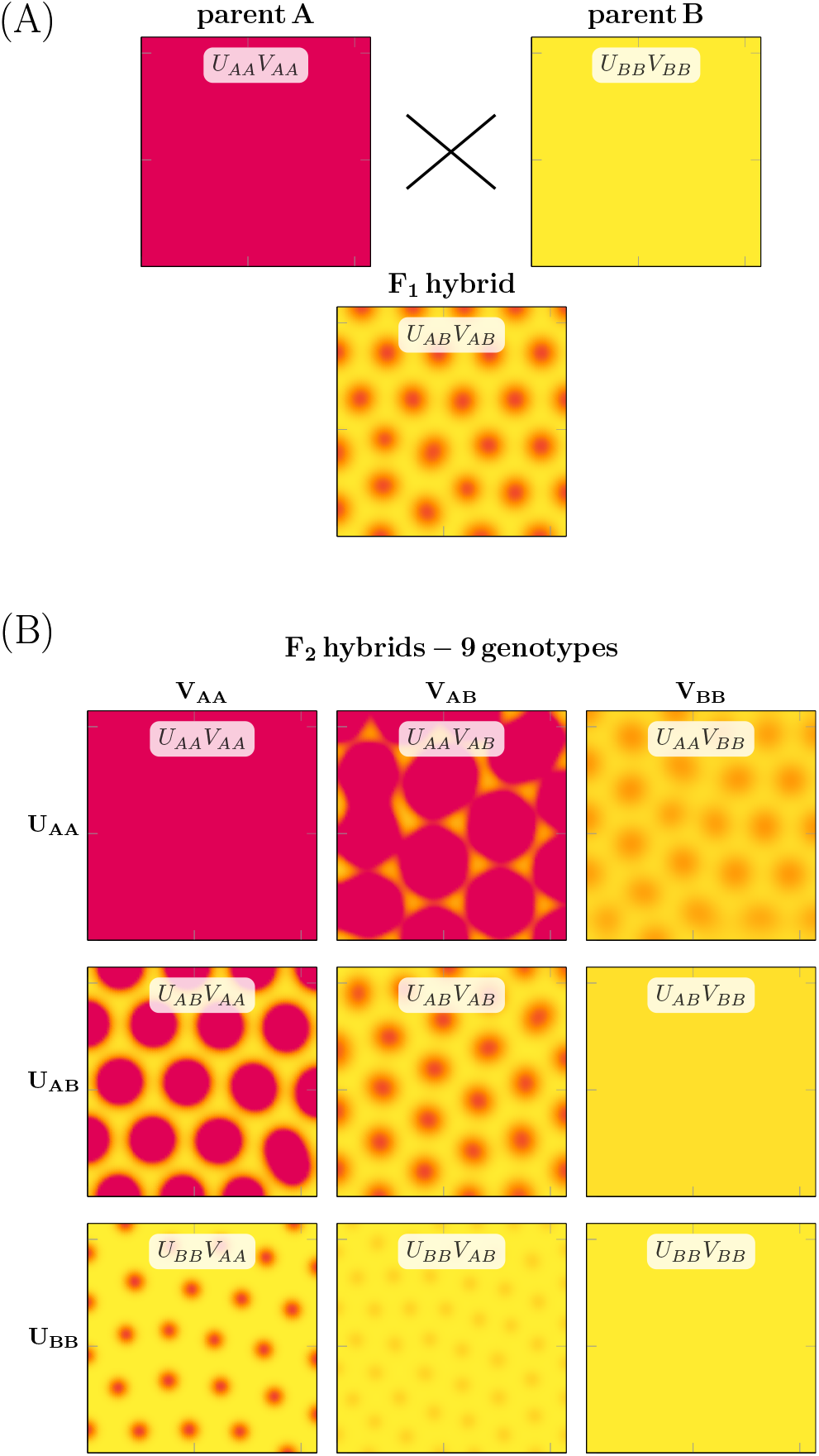
Multi-generational simulation of transgressive petal pattern phenotypes. (A) The diploid reaction-diffusion model (Eq. 4 and Fig. 3) produces unpatterned phenotypes when using homozygous parent parameters. Using heterozygous parameters corresponding to the cross between unpatterned parents (P_A_ *×* P_B_ *→* F1), the model produces patterned phenotypes in F1 offspring. Yellow-to-red pseudocolor represents the total activator concentration (i.e., low to high, *u* + *û*). (B) Simulated breeding of F1 hybrids (F1 *×* F1 *→* F2) yields various phenotypes, each corresponding to one of nine *F*_2_ genotypes (*U*_*AA*_*V*_*AA*_, *U*_*AB*_*V*_*AB*_, etc.). These diploid genotypes result from all possible combinations of parent haplotypes (denoted by subscripts *A* and *B*) that are composed of two alleles each for activator (*U*) and inhibitor (*V*) loci. Note: the F2 individuals with genotypes *U*_*AA*_*V*_*AA*_, *U*_*AB*_*V*_*AB*_, and *U*_*BB*_*V*_*BB*_ are identical to parent A, the *F*_1_ hybrid, and parent B, respectively. Parameters as in Table 4.

Fig. 4B shows the simulated pheonotypes of the F2 generation. The 3 *×* 3 grid organizes the results by genotype. The parent and hybrid F1 cases are recapitulated on the diagonal. As in the *Mimulus* experimental system, the simulated F2 hybrid population exhibits a wide variety of phenotypes. For this parameter set, 5 of the 6 phenotypes unique to the F2 generation are patterned (Turing unstable) while 1 is solid (Turing stable). The patterned phenotypes consist of spots with different intensity, size, and wavelength (e.g, spots in *U*_*BB*_*V*_*AA*_ and *U*_*BB*_*V*_*AB*_ are similar in size, but those in *U*_*BB*_*V*_*AB*_ are lighter and closer together).

**Table 4:** Parameters and critical values for Fig. 4. In all cases, *D*_*u*_ = 0.1.

|  |  |  |  |
| --- | --- | --- | --- |
| $\alpha_u^A = 2.8326 \times 10^{-5}$ | $\alpha_u^B = 9.5297 \times 10^{-4}$ | $\alpha_v^A = 2.4107 \times 10^{-4}$ | $\alpha_v^B = 0.0012$ |
| $\beta_u^A = 0.0018$ | $\beta_u^B = 0.0049$ | $\beta_v^A = 0.0067$ | $\beta_v^B = 0.0339$ |
| $\gamma_u^A = 1.1206$ | $\gamma_u^B = 0.8113$ | $\gamma_v^A = 0.0849$ | $\gamma_v^B = 0.3052$ |
| $\kappa_{uu}^{AA} = \kappa_{uu}^{AB} = 0.3575$ | $\kappa_{uu}^{BB} = \kappa_{uu}^{BA} = 0.2487$ | $\kappa_{vu}^{AA} = \kappa_{vu}^{AB} = 0.1589$ | $\kappa_{vu}^{BB} = \kappa_{vu}^{BA} = 0.1632$ |
| $\kappa_{uv}^{AA} = \kappa_{uv}^{AB} = 9.6994$ | $\kappa_{uv}^{BB} = \kappa_{uv}^{BA} = 7.5912$ | $\kappa_{vv}^{AA} = \kappa_{vv}^{AB} = 1.3040$ | $\kappa_{vv}^{BB} = \kappa_{vv}^{BA} = 0.2821$ |
| $d = 138.2$ | $d_\star^A = 142.3$ | $d_\star^B = 212.9$ | $d_\star^{F1} = 101.9$ |

Fig. 5 shows four more simulated F2 hybrid populations derived from unpatterned parents whose cross yields a patterned F1 hybrid (each displayed in a 3 *×* 3 grid akin to that in Fig. 4B). In three of the four examples (panels B, C, and D), the unique F2 phenotypes include 1 or more individuals with a pattern distinct from the F1 hybrid. The amount of phenotypic diversity among the F2 hybrids—the number of patterned individuals as well as the variety of patterns—can be low (panel A) or high (panel D) depending on model parameters.

**Figure 5:**
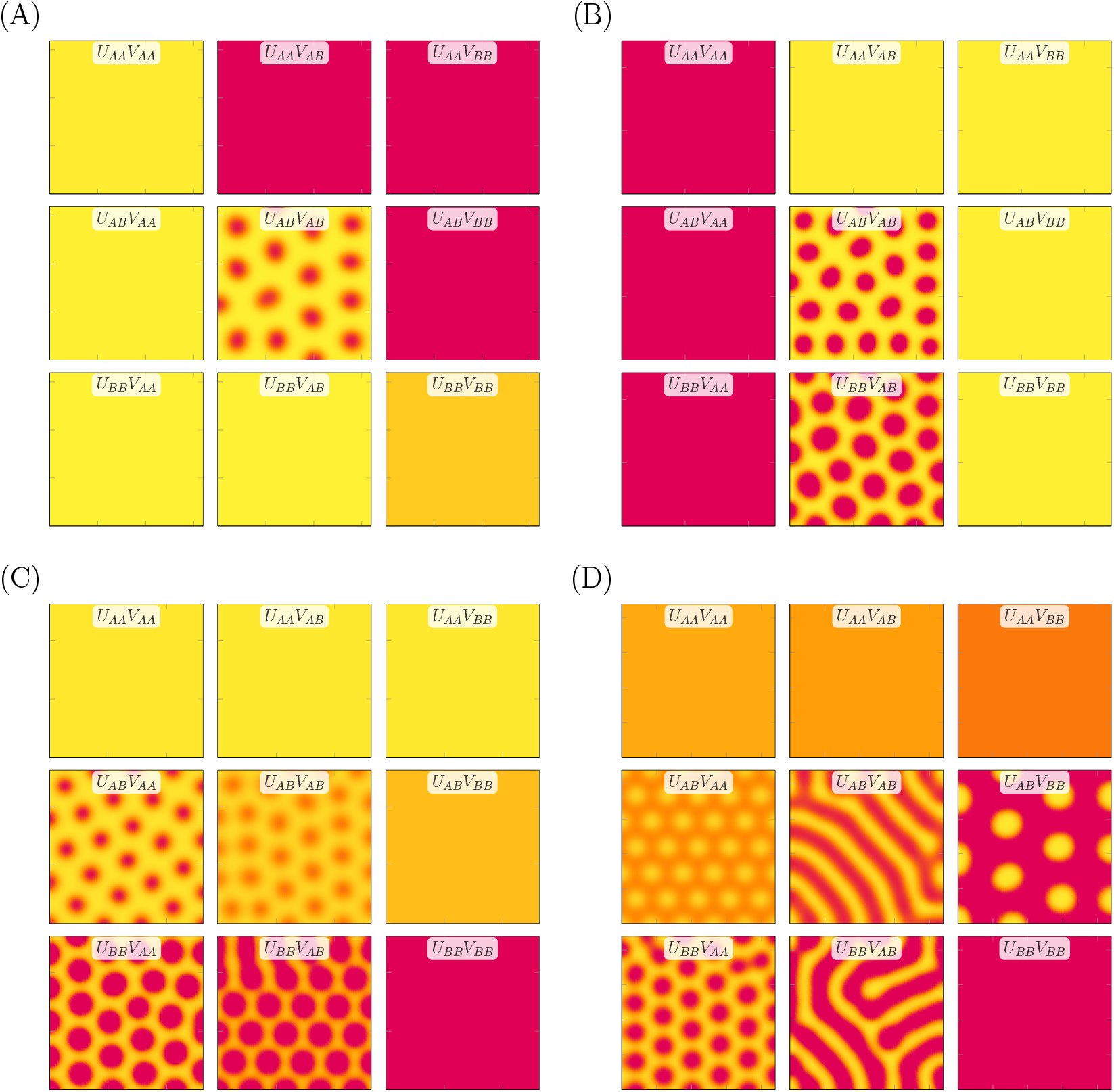
Four more examples of simulated selfing of transgressive patterned F1 hybrids. Panels (A)-(D) are ordered to show increasing pattern variety of the F2 hybrid populations. (A) No new patterns arise in the F2 generation. (B) One new F2 pattern is similar to the F1 hybrid. (C) Three new F2 patterns with spots larger and more intense than the F1 hybrid. (D) Four new F2 patterns, both labyrinthine and spotted phenotypes with various wavelengths, spot size, and spot intensity.

### 3.2 Parents with identical unpatterned phenotypes can produce patterned offspring

The multi-generational Turing model to produces a transgressive patterned F1 phenotype and a broad distribution of F2 phenotypes (Section 3.1), consistent with the experimental *Mimulus* system. Note that the cross between *M. l. variegatus* and *M. cupreus* (Fig. 1) involves unpatterned parents that are phenotypically distinct (*M. l. variegatus* is pink, *M. cupreus* is yellow). This is recapitulated in the simulated cross of Fig. 4, in which Parent A has a high concentration of activator (as does *M. l. variegatus*) while Parent B has a low activator concentration (like yellow *M. cupreus*).

Fig. 5A shows that our model formulation can produce transgressive F1 phenotypes even when the unpatterned parents are phenotypically similar (two shades of yellow). This observation highlights the transgressive nature of the patterned F1 and F2 phenotypes produced by the model. That is, the maxima of spatially inhomogeneous activator concentration in patterned F1 and F2 individuals can exceed the spatially homogenous activator concentrations of both unpatterned parents (see, e.g., genotype *U*_*AB*_*V*_*AB*_ in Fig. 6A). In some cases, unpatterned F2 individuals exhibit activator concentration that is more extreme than either parent (*U*_*AA*_*V*_*BB*_ in Fig. 6A).

**Figure 6:**
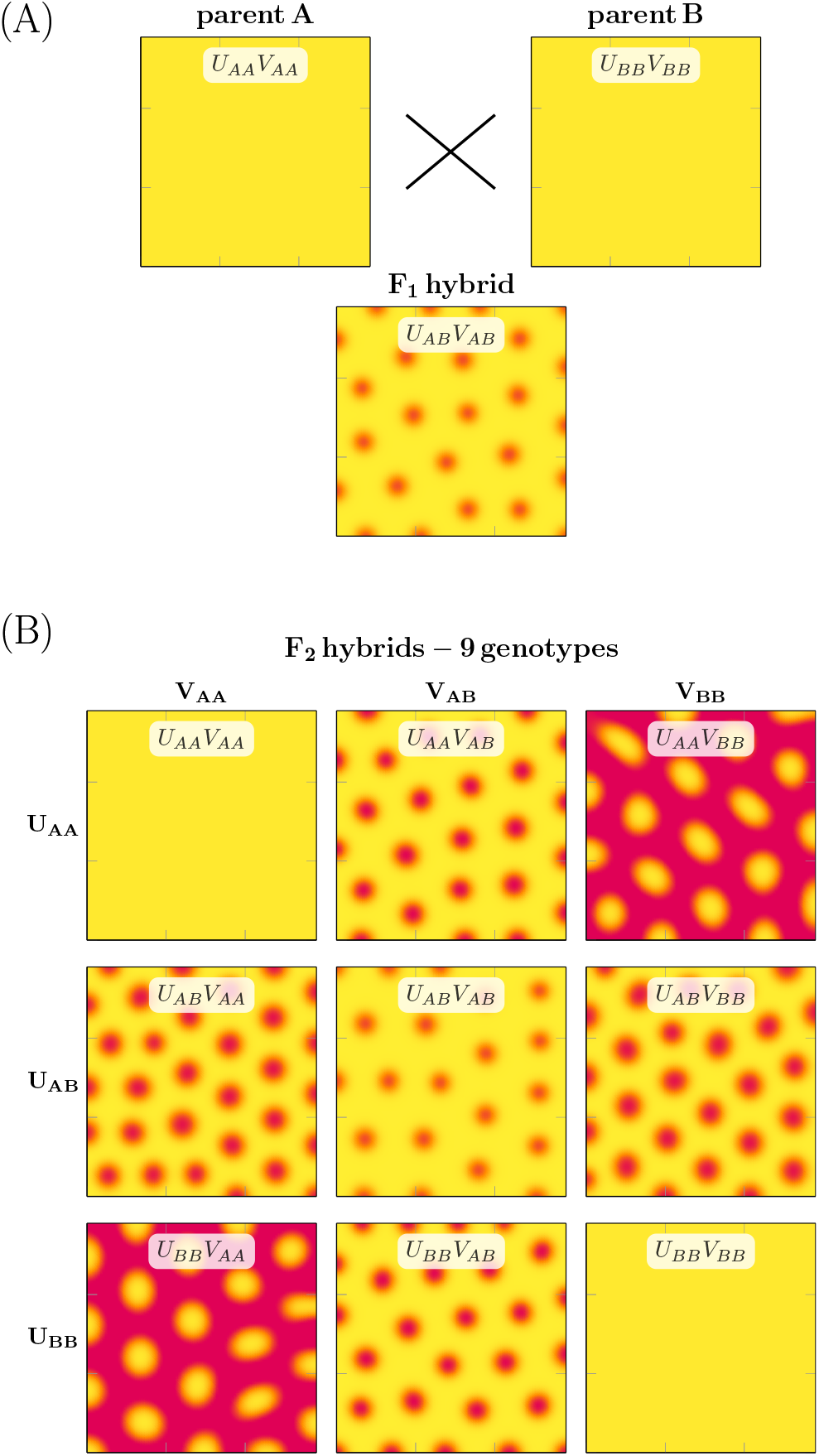
Transgressive patterns arising from phenotypically identical parents. (A) The diploid reaction-diffusion model (Eq. 4) can produce transgressive patterns in the F1 generation even when the homozygous parents have identical unpatterned phenotypes. In the simulated cross (*P*_*A*_ *× P*_*B*_ *→ F*_1_), the heterozygous parameter set is identical to the parents except for trans-interaction binding affinities (e.g., 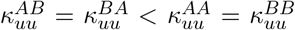, and similarly for *κ*_*uv*_, *κ*_*vu*_, and *κ*_*vv*_). (B) The *F*_2_ hybrids include five distinct phenotypes, four of which are patterned.

### 3.3 A possible mechanism for the emergence of transgressive F1 hybrid phenotypes

Fig. 5A shows a transgressive F1 hybrid (red spots on yellow background) from a cross between phenotypically similar unpatterned parents (one yellow and one light orange). Interestingly, Fig. 6 shows that phenotypically *identical* unpatterned parents (*U*_*AA*_*V*_*AA*_ and *U*_*BB*_*V*_*BB*_) can yield patterned F1 hybrid offspring (*U*_*AB*_*V*_*AB*_) and novel patterned F2 phenotypes. We will discuss this limit case in detail because it suggests one possible mechanism for the emergence of transgressive phenotypes in hybrid *Mimulus*.

The parents in Fig. 6 are phenotypically identical (unpatterned yellow) because, in the parameter set used, the rate constants do not depend on the allele type (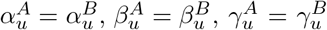, and similarly for *v*). The eight binding constants used in the parent simulations (both doubly homozygous genotypes) also do not depend on allele 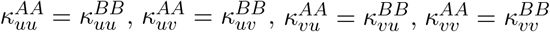. For parameters with this symmetry, the reaction-diffusion systems representing parent A and B are the same; consequently, the parent simulations yield identical phenotypes (unpatterned yellow, see *U*_*AA*_*V*_*AA*_ and *U*_*BB*_*V*_*BB*_ in Fig. 6). The simulation of the doubly heterozygous F1 hybrid involves these parental parameters as well as eight additional binding constants chosen to satisfy 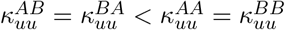 (and similarly for *κ*_*uv*_, *κ*_*vu*_, and *κ*_*vv*_). This choice is consistent with small structural differences in the transcription factors from alleles A and B leading to less effective binding to the regulatory side associated with the alternate allele type.

The central panel of Fig. 6 (*U*_*AB*_*V*_*AB*_) shows that a patterned F1 hybrid is possible under the assumptions made in the previous paragraph. In this simulation, the F2 hybrids include five distinct phenotypes, four of which are patterned. Each of the three patterned phenotypes that are unique to the F2 population is produced by two equivalent genotypes (*U*_*AA*_*V*_*AB*_ = *U*_*BB*_*V*_*AB*_, *U*_*AB*_*V*_*AA*_ = *U*_*AB*_*V*_*BB*_, *U*_*AA*_*V*_*BB*_ = *U*_*BB*_*V*_*AA*_). Note that these genotype pairs result from swapping one or both of the homozygous loci (activator *U*_*AA*_ *↔ U*_*BB*_ or inhibitor *V*_*AA*_ *↔ V*_*BB*_). For example, replacing *U*_*AA*_ by *U*_*BB*_ in *U*_*AA*_*V*_*AB*_ gives *U*_*BB*_*V*_*AB*_.

Fig. 6 demonstrates that transgressive phenotypes in doubly heterozygous F1 hybrids can emerge when the TFs derived from one parent are less effective at regulating TFs derived from the other parent (with different allele type). For example, the inequality 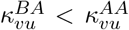 means that the activator of allele type A has lower affinity for the binding site of inhibitor B than A. We will refer to regulation between identical and distinct allele types as *cis* and *trans*, respectively. In *cis* regulation, both the transcription factor and its regulatory binding site are derived from the same allele type. In *trans* regulation, the transcription factor regulates the production of the alternate allele’s gene product. To explore this mechanism further, observe that the parameter symmetries of this limit case allow the reaction-diffusion equations for the F1 hybrid to be contracted. Writing the total concentration of activator and inhibitor as 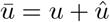 and 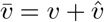, Eq. 6 is equivalent to

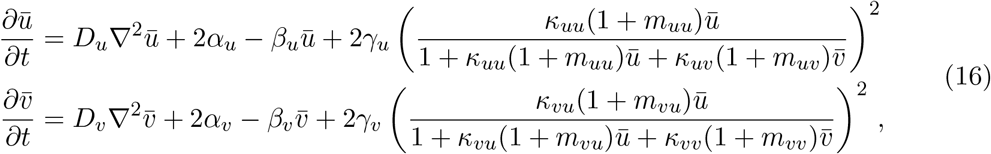

where the superscripted A and B are dropped because 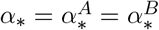 (similarly for *β* and *γ*) and 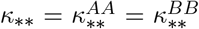. In this equation, *m*_*\*\**_ is the ratio of *trans* to *cis* binding constants, 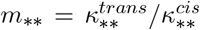, where 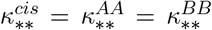 and 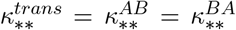 represent affinities within and between parental haplotypes, respectively. The assumption that *trans* binding is less effective than *cis* binding implies that 0 *≤ m*_*\*\**_ *<* 1. Setting *m*_*\*\**_ = 1 gives equations for the phenotypically identical parents.

Fig. 7 summarizes a parameter study for which the only distinction between parents and F1 hybrid parameters is a decreased *trans* binding affinity for one of the four regulatory pathways (*κ*_*uu*_, *κ*_*uv*_, *κ*_*vu*_, or *κ*_*vv*_). Fig. 7A shows that reduction of the trans efficacy of the activator regulating inhibitor (decreasing *m*_*vu*_) decreases the critical diffusion ratio *d*_*⋆*_. This suggests the following parsimonious hypothesis for the emergence of transgressive phenotypes in an F1 hybrid derived from phenotypically similar parents—a Turing bifurcation occurs as a result of a decrease in the inhibitor production rate, 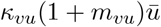, which is 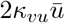 in the parents, but a lesser value in the hybrid (between 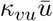 and 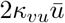). Fig. 7C shows that a similar result is obtained with a reduction of the trans efficacy of the inhibitor binding the activatory regulatory site (decreasing *m*_*uv*_).

**Figure 7:**
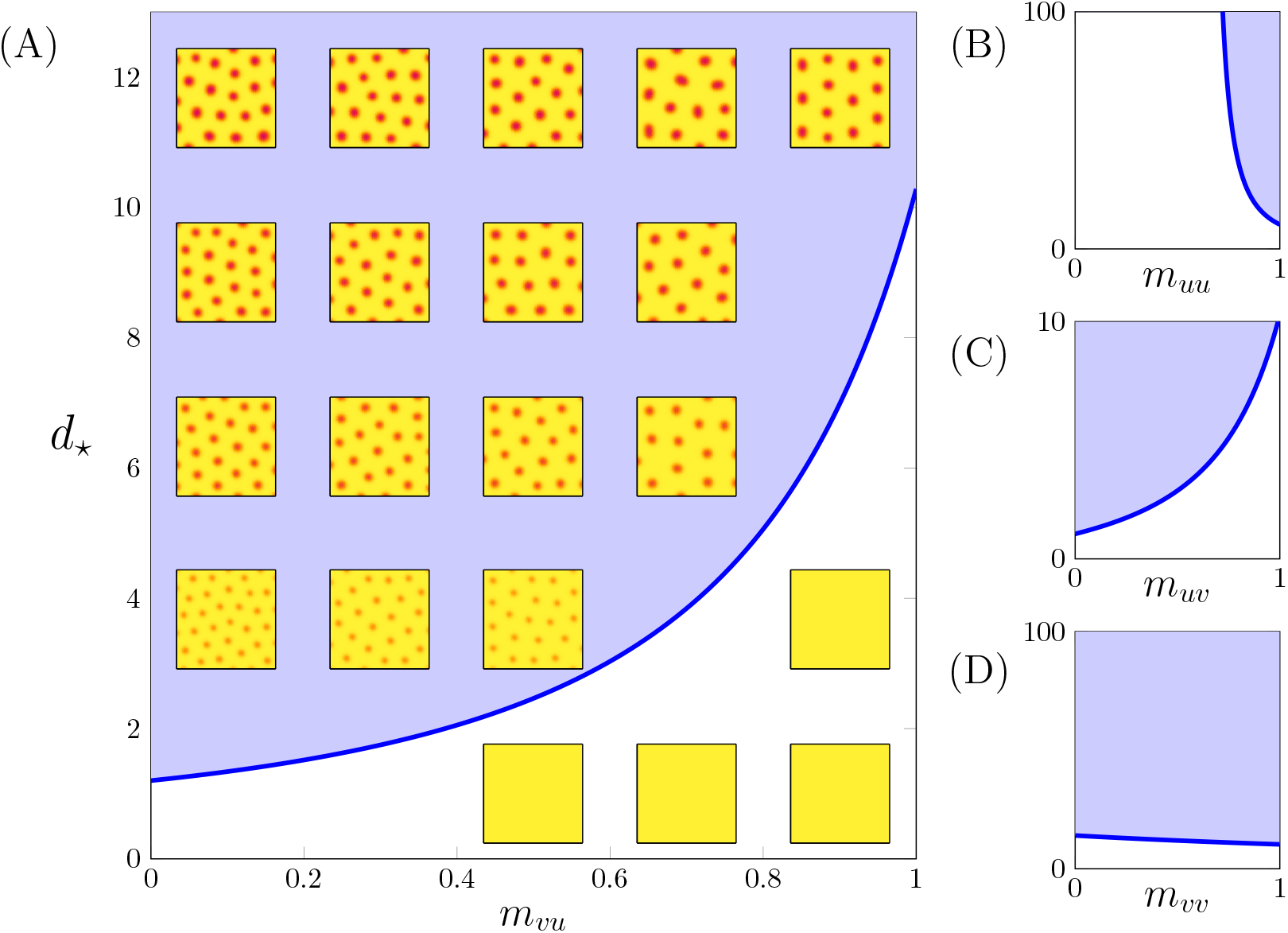
Reducing the trans-interaction efficacy affects the critical diffusion ratio *d*_*⋆*_. The trans efficacy of transcription factor binding is the ratio of two affinities (*m*_*uu*_, *m*_*uv*_, *m*_*vu*_, and *m*_*vv*_). In panel (A) the parameter *m*_*vu*_ (horizontal axis) is defined as 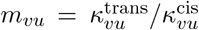 where 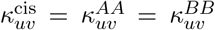 and 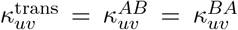 represent affinities within and between parental haplotypes, respectively. The critical diffusion coefficient ratio (*d*_*⋆*_ = *D*_*v*_*/D*_*u*_, vertical axis) locates a Turing instability (i.e., spotted patterns occur above the blue curve). Reducing the trans efficacy *m*_*vu*_ decreases *d*_*⋆*_. Reducing the trans efficacies *m*_*uu*_ (B), *m*_*uv*_ (C), and *m*_*vv*_ (D) impact *d*_*⋆*_ with different directions and intensities. Parameters as in Fig. 6.

Fig. 7 demonstrates that reduction in trans efficacy of inhibitor regulating activator (*m*_*uv*_) or activator regulating inhibitor (*m*_*vu*_) can lead to instability in the F1 hybrid, but this is not the case for *m*_*uu*_ or *m*_*vv*_. That is, decreasing *m*_*uv*_ (or *m*_*vu*_) decreases *d*_*⋆*_, but decreasing *m*_*uu*_ (or *m*_*vv*_) increases *d*_*⋆*_. These results depend on the parameters chosen for the phenotypically identical parents (Eq. 16 with *m*_*\*\**_ = 1). Nevertheless, Fig. 7 is representative of the most common outcomes we have observed for admissible parameter sets (Turing stable parents, Turing unstable hybrid).

## 4 Discussion

This paper presents a multi-generational Turing model of pattern formation in hybrid *Mimulus*. Model development was motivated by experimental observations of petal phenotypes across three generations: parents, F1, and F2 (Fig. 1). Our model formulation is explicitly diploid, i.e., there is a representation of multiple copies of each genetic loci—two for activator, two for inhibitor (Fig. 3). The model reproduces transgressive phenotypes in F1 hybrids between two unpatterned parents, i.e., red *×* yellow *→* red spots on yellow (Fig. 4). Consistent with experiments, simulated selfing of this F1 hybrid often yields a distribution of phenotypes in the F2 hybrid population (Fig. 5).

It is instructive to compare our model with the inheritance of an activator-inhibitor reaction-diffusion system motivated by pattern formation in zebrafish [Miyazawa et al., 2010]. This prior work, which is the only other example of a multi-generational Turing model to be found in the literature, presumes that zebrafish phenotypes are controlled by a large number of genetic loci. Assuming additive inheritance, model F1 hybrids using parameter values intermediate to parental values. This approach generated novel phenotypes in F1 hybrids. However, the F1 hybrid retains no information regarding the more extreme parental parameter values; it is unable to reproduce a distribution of phenotypes in an F2 population. In contrast, the multi-generational model presented here allows for loci to be homozygous or heterozygous for parental allele type. Simulated selfing of F1 hybrids leads to 9 distinct F2 genotypes and a wide range of phenotypes, consistent with monkeyflower experiments (Fig. 5). This is the preferred approach for a model of the anthocyanin pathway in *Mimulus*, which is known to be controlled by a small number of genes.

Under the model put forth by Miyazawa et al. [2010], offspring phenotype will necessarily be intermediate to parental phenotype. However, it is well documented that hybridization can lead to offspring phenotype that is more extreme than either parent (see Birchler et al., 2010; Hochholdinger and Baldauf, 2018, for reviews of hybrid vigor and heterosis). We find that our explicitly diploid model can yield offspring with phenotypes more extreme than either parent (Fig. 5A). As a limiting case, we performed simulations in which parents are phenotypically identical. These simulations suggest a hypothetical mechanism for the emergence of hybrid phenotypes involving transcription factors whose regulatory binding is weaker between parental alleles versus within parental alleles. In principle, this hypothesis could be empirically tested using ChIP-Seq to identify binding sites and relative strengths of binding (see Liang and Kele, 2012, for details).

Although there is experimental evidence for an activator-inhibitor reaction-diffusion system in *Mimulus* Ding et al., 2020, the anthocyanin pathway is known to include two accessory proteins, bHLH and WD40, and at least two copies of the inhibitor RTO that may bind and sequester bHLH [Ding et al., 2020; Yuan et al., 2014]. In future work, this paper’s approach to diploidy and inheritance could be used to study patterning mediated by a gene regulatory network model with these realistic features.

This paper is a proof of concept that emphasizes a specific biological question, model formulation, and numerical simulations. Each multi-generational simulation presented here is, in fact, nine distinct reaction-diffusion systems and solutions and their numerically calculated steady states. These systems are related to one another in a complex manner through combinations of parameter assignments that account for Mendelian inheritance of multiple alleles and gene products. Because there are no continuously changing (bifurcation) parameters in our representation of inheritance, it is not clear (to us) what form a deeper mathematical analysis would take.

## Appendix A: Equations and contractions for F2 genotypes

In the equations for the diploid F2 population, the four-variable system can be contracted by one equation per instance of homozygosity. For the parent with homozygous genotype *U*_*AA*_*V*_*AA*_, the four-variable system (Eq. 7) can be contracted to a two-variable system (Eq. 8). The equations used for parent B (*U*_*BB*_*V*_*BB*_) are identical with the replacement of *B* for *A*. In contrast, simulations of the doubly heterozygous F1 hybrid with genotype *U*_*AB*_*V*_*AB*_ require all four equations (Eq. 6). Six additional genotypes are unique to the F2 population. The equations for these genotypes (contracted when possible) are enumerated below.

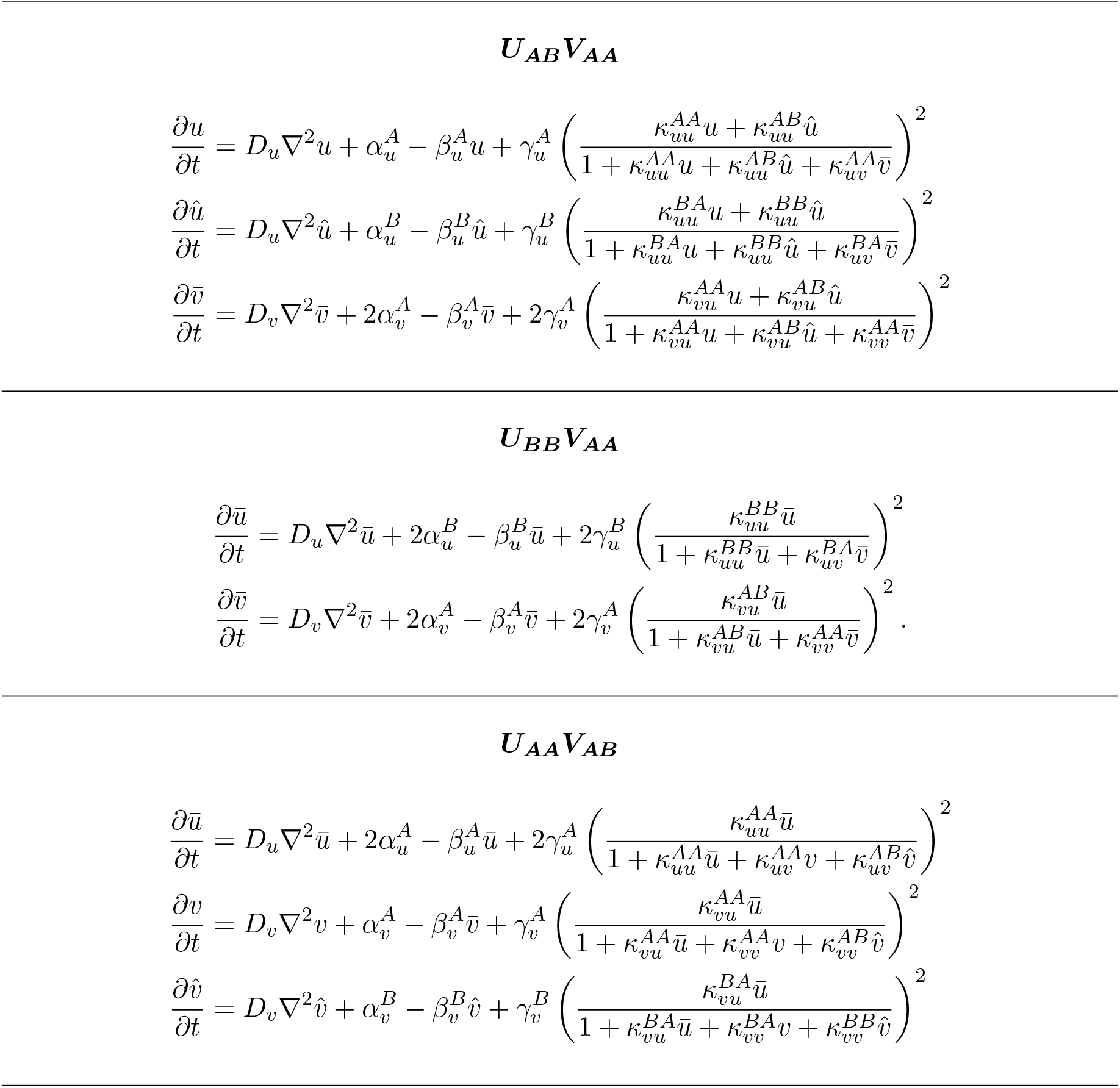
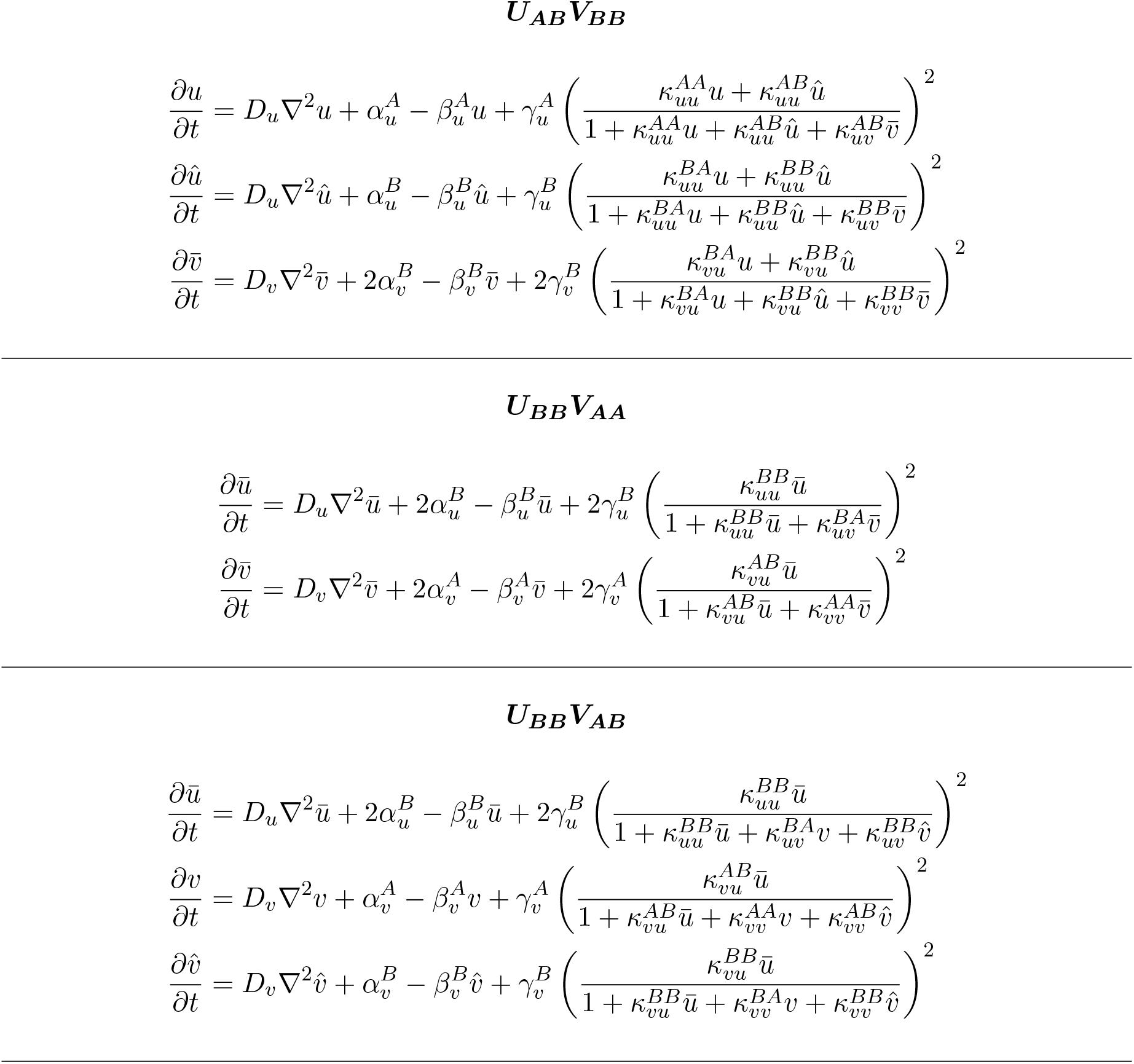

## Appendix B: Selection of multi-generational parameter sets

The parameters of Figs. 4–6 were chosen as follows. First, rate and binding constants needed for an F1 hybrid were selected as i.i.d. exponentially distributed random variables with prescribed means 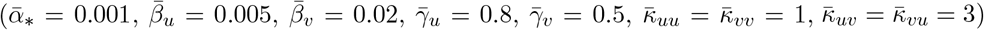. Next, the appropriate subsets of parameters were assigned to parents A and B as outlined in Section 2. A random parameter set was considered admissible if Parents A and B were Turing stable, while the F1 hybrid is Turing unstable (see Section 2.4). Stability of the *J* and *J* ^*′*^ (Eqs. 12–15) was ascertained by numerical calculation of eigenvalues. This requires the steady state of the spatially homogeneous solution for each individual, obtained by numerically integrating the appropriate ODE system of reaction terms. Finally, we fixed *D*_*u*_ and calculated (using a binary search) the critical values of *D*_*v*_ = *dD*_*u*_ for parent A, parent B, and F1 hybrid leading to unstable *J* ^*′*^. If the corresponding critical diffusion coefficient ratios satisfied 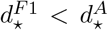 and 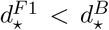, the multi-generational parameter was considered admissible, because choosing *D*_*v*_ so that 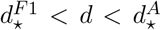 and 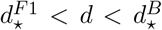 yields Turing-stable parents and a Turing-unstable F1 hybrid (i.e., a transgressive phenotype).

Tables 4 and 5 record the parameters used for Figs. 4 and 5. Because the doubly heterozygous F1 hybrid simulation uses every parameter, and all other genotypes use a subset of these, it suffices to tabulate the F1 hybrid parameters (recall Section 2.2, Tables 1 and 2). In Tables 4 and 5, the degradation rates (*β*_*\**_) are first-order rate constants with physical dimensions of inverse time (1/time). The production rates (*α*_*\**_ and *γ*_*\**_) are zeroth order (concentration/time). The binding constants *κ*_*\*\**_ are equilibrium association constants with physical dimensions of inverse concentration (1/concentration). Because all calculations are steady states, time can be considered dimensionless, and rates are meaningful primarily through their relative values. The spatial extent of the reaction-diffusion calculations is 2–20 millimeters (width and height).

**Table 5:** Parameters used in Fig. 5. In all cases, *D*_*u*_ = 0.1.

| Fig. 5A |  |  |  |
| --- | --- | --- | --- |
| $\alpha_u^A = 5.0617 \times 10^{-4}$ | $\alpha_u^B = 4.8463 \times 10^{-5}$ | $\alpha_v^A = 7.5301 \times 10^{-4}$ | $\alpha_v^B = 6.3299 \times 10^{-4}$ |
| $\beta_u^A = 0.0048$ | $\beta_u^B = 0.0029$ | $\beta_v^A = 0.0094$ | $\beta_v^B = 0.0182$ |
| $\gamma_u^A = 1.1298$ | $\gamma_u^B = 0.8056$ | $\gamma_v^A = 1.2327$ | $\gamma_v^B = 0.4460$ |
| $\kappa_{uu}^{AA} = \kappa_{uu}^{AB} = 3.9125$ | $\kappa_{uu}^{BB} = \kappa_{uu}^{BA} = 1.5698$ | $\kappa_{vu}^{AA} = \kappa_{vu}^{AB} = 6.4121$ | $\kappa_{vu}^{BB} = \kappa_{vu}^{BA} = 0.2480$ |
| $\kappa_{uv}^{AA} = \kappa_{uv}^{AB} = 3.1422$ | $\kappa_{uv}^{BB} = \kappa_{uv}^{BA} = 15.0419$ | $\kappa_{vv}^{AA} = \kappa_{vv}^{AB} = 0.4017$ | $\kappa_{vv}^{BB} = \kappa_{vv}^{BA} = 0.4074$ |
| $d = 40.4$ | $d_\star^A = 41.3$ | $d_\star^B = 51.5$ | $d_\star^{F1} = 31.3$ |
| Fig. 5B |  |  |  |
| $\alpha_u^A = 0.0016$ | $\alpha_u^B = 1.0142 \times 10^{-4}$ | $\alpha_v^A = 7.4401 \times 10^{-4}$ | $\alpha_v^B = 1.7396 \times 10^{-4}$ |
| $\beta_u^A = 0.0052$ | $\beta_u^B = 0.0022$ | $\beta_v^A = 0.0248$ | $\beta_v^B = 0.0131$ |
| $\gamma_u^A = 1.7306$ | $\gamma_u^B = 0.4177$ | $\gamma_v^A = 0.8788$ | $\gamma_v^B = 0.8070$ |
| $\kappa_{uu}^{AA} = \kappa_{uu}^{AB} = 0.5348$ | $\kappa_{uu}^{BB} = \kappa_{uu}^{BA} = 0.4988$ | $\kappa_{vu}^{AA} = \kappa_{vu}^{AB} = 1.5258$ | $\kappa_{vu}^{BB} = \kappa_{vu}^{BA} = 4.7691$ |
| $\kappa_{uv}^{AA} = \kappa_{uv}^{AB} = 2.4156$ | $\kappa_{uv}^{BB} = \kappa_{uv}^{BA} = 1.0868$ | $\kappa_{vv}^{AA} = \kappa_{vv}^{AB} = 1.9042$ | $\kappa_{vv}^{BB} = \kappa_{vv}^{BA} = 1.0809$ |
| $d = 180.6$ | $d_\star^A = 243.2$ | $d_\star^B = 182.6$ | $d_\star^{F1} = 162.5$ |
| Fig. 5C |  |  |  |
| $\alpha_u^A = 1.1154 \times 10^{-4}$ | $\alpha_u^B = 2.9340 \times 10^{-4}$ | $\alpha_v^A = 1.7720 \times 10^{-4}$ | $\alpha_v^B = 1.4803 \times 10^{-4}$ |
| $\beta_u^A = 0.0067$ | $\beta_u^B = 0.0071$ | $\beta_v^A = 0.0070$ | $\beta_v^B = 0.0154$ |
| $\gamma_u^A = 0.1384$ | $\gamma_u^B = 0.1528$ | $\gamma_v^A = 0.5847$ | $\gamma_v^B = 0.4356$ |
| $\kappa_{uu}^{AA} = \kappa_{uu}^{AB} = 1.1426$ | $\kappa_{uu}^{BB} = \kappa_{uu}^{BA} = 1.7916$ | $\kappa_{vu}^{AA} = \kappa_{vu}^{AB} = 0.4669$ | $\kappa_{vu}^{BB} = \kappa_{vu}^{BA} = 1.4204$ |
| $\kappa_{uv}^{AA} = \kappa_{uv}^{AB} = 9.1243$ | $\kappa_{uv}^{BB} = \kappa_{uv}^{BA} = 4.2542$ | $\kappa_{vv}^{AA} = \kappa_{vv}^{AB} = 1.3774$ | $\kappa_{vv}^{BB} = \kappa_{vv}^{BA} = 2.6028$ |
| $d = 40.4$ | $d_\star^A = 535.8$ | $d_\star^B = 41.4$ | $d_\star^{F1} = 31.3$ |
| Fig. 5D |  |  |  |
| $\alpha_u^A = 0.0011$ | $\alpha_u^B = 5.7845 \times 10^{-4}$ | $\alpha_v^A = 3.9353 \times 10^{-4}$ | $\alpha_v^B = 0.0024$ |
| $\beta_u^A = 0.0078$ | $\beta_u^B = 0.0039$ | $\beta_v^A = 0.0512$ | $\beta_v^B = 0.0072$ |
| $\gamma_u^A = 0.0873$ | $\gamma_u^B = 0.1341$ | $\gamma_v^A = 1.3311$ | $\gamma_v^B = 0.0466$ |
| $\kappa_{uu}^{AA} = \kappa_{uu}^{AB} = 2.4419$ | $\kappa_{uu}^{BB} = \kappa_{uu}^{BA} = 1.3145$ | $\kappa_{vu}^{AA} = \kappa_{vu}^{AB} = 0.8460$ | $\kappa_{vu}^{BB} = \kappa_{vu}^{BA} = 2.8702$ |
| $\kappa_{uv}^{AA} = \kappa_{uv}^{AB} = 2.5520$ | $\kappa_{uv}^{BB} = \kappa_{uv}^{BA} = 2.7183$ | $\kappa_{vv}^{AA} = \kappa_{vv}^{AB} = 0.8983$ | $\kappa_{vv}^{BB} = \kappa_{vv}^{BA} = 1.5572$ |
| $d = 575.2$ | $d_\star^A = 878.9$ | $d_\star^B = 596.4$ | $d_\star^{F1} = 384.5$ |

**Table 6:**
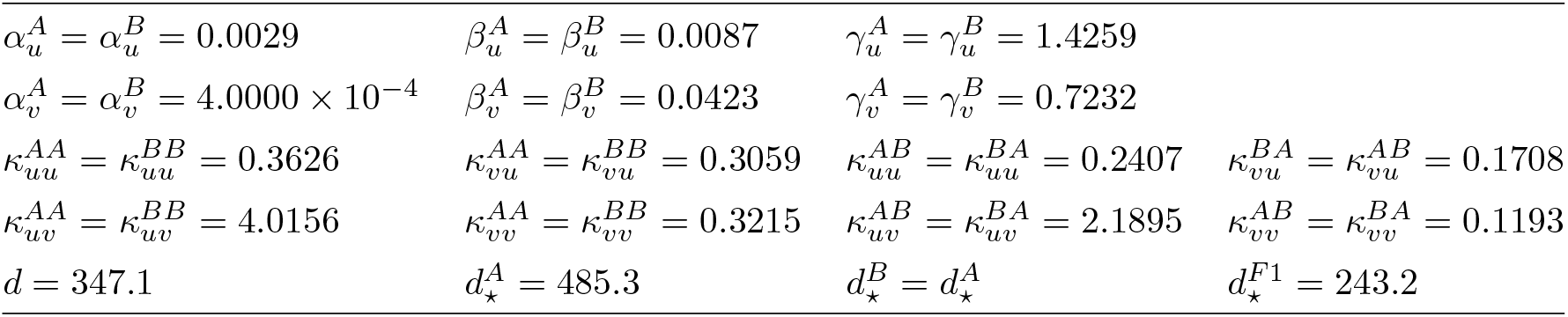
Parameters used in Fig. 6. Parents have equal rate constants (e.g., 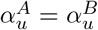) and binding constants 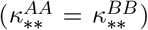. The F1 hybrid has reduced trans-interaction binding coefficients, that is, 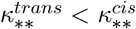 where 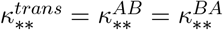 and 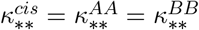.

|  |  |  |  |
| --- | --- | --- | --- |
| $\alpha_u^A = \alpha_u^B = 0.0029$ | $\beta_u^A = \beta_u^B = 0.0087$ | $\gamma_u^A = \gamma_u^B = 1.4259$ | |
| $\alpha_v^A = \alpha_v^B = 4.0000 \times 10^{-4}$ | $\beta_v^A = \beta_v^B = 0.0423$ | $\gamma_v^A = \gamma_v^B = 0.7232$ | |
| $\kappa_{uu}^{AA} = \kappa_{uu}^{BB} = 0.3626$ | $\kappa_{vu}^{AA} = \kappa_{vu}^{BB} = 0.3059$ | $\kappa_{uu}^{AB} = \kappa_{uu}^{BA} = 0.2407$ | $\kappa_{vu}^{BA} = \kappa_{vu}^{AB} = 0.1708$ |
| $\kappa_{uv}^{AA} = \kappa_{uv}^{BB} = 4.0156$ | $\kappa_{vv}^{AA} = \kappa_{vv}^{BB} = 0.3215$ | $\kappa_{uv}^{AB} = \kappa_{uv}^{BA} = 2.1895$ | $\kappa_{vv}^{AB} = \kappa_{vv}^{BA} = 0.1193$ |
| $d = 347.1$ | $d_{\star}^A = 485.3$ | $d_{\star}^B = d_{\star}^A$ | $d_{\star}^{F1} = 243.2$ |

## Acknowledgements

The work was supported in part by National Science Foundation grants DEB #2031275 to AMC, JRP & GDCS, and DMS #1951646 to GDCS. ESGS was jointly mentored by JRP & GDCS.

## References

N. W. Albert, K. M. Davies, D. H. Lewis, H. Zhang, M. Montefiori, C. Brendolise, M. R. Boase, H. Ngo, P. E. Jameson, and K. E. Schwinn. A conserved network of transcriptional activators and repressors regulates anthocyanin pigmentation in eudicots. The Plant Cell, 26(3):962–980, 2014. doi: 10.1105/tpc.113.122069.

J. A. Birchler, H. Yao, S. Chudalayandi, D. Vaiman, and R. A. Veitia. Heterosis. The Plant Cell, 22(7):2105–2112, 2010. doi: 10.1105/tpc.110.076133.

D. Bouyer, F. Geier, F. Kragler, A. Schnittger, M. Pesch, K. Wester, R. Balkunde, J. Timmer, C. Fleck, and M. Hülskamp. Two-dimensional patterning by a trapping/depletion mechanism: The role of TTG1 and GL3 in Arabidopsis trichome formation. PLoS Biology, 6(6):e141, 2008. doi: 10.1371/journal.pbio.0060141.

M. Cooley and J. H. Willis. Genetic divergence causes parallel evolution of flower color in Chilean Mimulus. New Phytologist, 183(3):729–739, 2009. doi: 10.1111/j.1469-8137.2009.02858.x.

Ding, E. L. Patterson, S. V. Holalu, J. Li, G. A. Johnson, L. E. Stanley, A. B. Greenlee, F. Peng, H. Bradshaw, M. L. Blinov, B. K. Blackman, and Y.-W. Yuan. Two MYB proteins in a self-organizing activator-inhibitor system produce spotted pigmentation patterns. Current Biology, 30(5):802–814, 2020. doi: https://doi.org/10.1016/j.cub.2019.12.067.

F. Hochholdinger and J. A. Baldauf. Heterosis in plants. Current Biology, 28(18):R1089– R1092, 2018.

T. Ishida, T. Kurata, K. Okada, and T. Wada. A genetic regulatory network in the development of trichomes and root hairs. Annual Review of Plant Biology, 59(1):365–386, 2008. doi: 10.1146/annurev.arplant.59.032607.092949.

T. Kurata, K. Okada, and T. Wada. Intercellular movement of transcription factors. Journal of Cell Biology, 8(6):600–605, 2005. doi: 10.1016/j.pbi.2005.09.005.

J. Larkin, N. Young, M. Prigge, and M. Marks. The control of trichome spacing and number in Arabidopsis. Development, 122(3):997–1005, 1996. doi: 10.1242/dev.122.3.997.

K. Liang and S. Kele. Detecting differential binding of transcription factors with ChIP-seq. Bioinformatics, 28(1):121–122, 2012.

H. Meinhardt. Turing’s theory of morphogenesis of 1952 and the subsequent discovery of the crucial role of local self-enhancement and long-range inhibition. Interface Focus, 2(4): 407–416, 2012. doi: 10.1098/rsfs.2011.0097.

S. Miyazawa, M. Okamoto, and S. Kondo. Blending of animal colour patterns by hybridization. Nature Communications, 1(1):66, 2010. doi: 10.1038/ncomms1071.

J. D. Murray. Mathematical Biology II: Spatial Models and Biomedical Applications, volume 3. Springer, 2001.

M. Pesch and M. Hülskamp. Creating a two-dimensional pattern de novo during Arabidopsis trichome and root hair initiation. Current Opinion in Genetics & Development, 14(4): 422–427, 2004. doi: 10.1016/j.gde.2004.06.007.

R. Stracke, M. Werber, and B. Weisshaar. The R2R3-MYB gene family in Arabidopsis thaliana. Current Opinion in Plant Biology, 4(5):447–456, 2001.

A. M. Turing. The chemical basis of morphogenesis. Bulletin of Mathematical Biology, 52 (1-2):153–197, 1990.

Y.-W. Yuan, J. M. Sagawa, L. Frost, J. P. Vela, and H. D. Bradshaw. Transcriptional control of floral anthocyanin pigmentation in monkeyflowers (Mimulus). New Phytologist, 204(4): 1013–1027, 2014. doi: 10.1111/nph.12968.

